# Deep Learning Behavioral Phenotyping System in the Diagnosis of Parkinson’s Disease with Drosophila melanogaster

**DOI:** 10.1101/2024.02.23.581846

**Authors:** Keyi Dong, April Burch, Kang Huang

**Author notes:** Corresponding Author: Dr. Kang Huang.

## Abstract

*Drosophila Melanogaster* is widely used as animal models for Parkinson’s disease (PD) research. Because of the complexity of MoCap and quantitative assessment among *Drosophila Melanogaster*, however, there is a technical issue that identify PD symptoms within drosophila based on objective spontaneous behavioral characteristics. Here, we developed a deep learning framework generated from kinematic features of body posture and motion between wildtype and SNCA^E46K^ mutant drosophila genetically modeled □-Syn, supporting clustering and classification of PD individuals. We record locomotor activity in a 3D-printed trap, and utilize the pre-analysis pose estimation software DeepLabCut (DLC) to calculate and generate numerical data representing the motion speed, tremor frequency, and limb motion of *Drosophila Melanogaster*. By plugging these data as the input, the diagnosis result (1/0) representing PD or WT as the output. Our result provides a toolbox which would be valuable in the investigation of PD progressing and pharmacotherapeutic drug development.

## Introduction

Parkinson’s disease (PD) is a recognizable neurodegeneration with complex etiology and rapid progress. Compared with the 1990s, the age standardized prevalence rate of PD increased by 21.7% over the same period^1^. Considerable evidence in recent studies showed that the progressive degeneration of dopaminergic (DA) neurons in the substantia nigra (SN) and the parenchyma (SNpc) was the main cause of the motor symptoms^2^, and the pathological hallmarks of PD are the deposition of Lewy bodies (LBs), which are mainly composed of misfolding α-synuclein (α-Syn)^3^. For this reason, several trans-genetic *Drosophila melanogaster*, such as E46K mutation of SNCA gene, have been utilized as animal models to explore the pathological mechanism of PD induced by single gene mutation^4-6^.

Compared with mammalian models such as rodents and NHPs (non-human primates), PD related trans-genetic *Drosophila melanogaster* has obvious advantages in distinguished physical signs, abnormal phenotypes in motor and non-motor behaviors, and short lifespan^7^. Whereas, due to the immatureness of the hierarchical and classified observation technology of entomological behaviors with *Drosophila melanogaster*, researchers still need a quantitative and time-scale dynamic phenotypic classification of *Drosophila melanogaster* during exploring the mechanism of PD affected by multiple factors, especially for non-transgenetic models^8^. Recently, scientists have developed and optimized a variety of software toolbox to track and quantify the behavioral characteristics of small but fast-moving animals including *Drosophila melanogaster*, such as DeepGraphPose, DeepPoseKit, DeepLabCut, LEAP^9-12^ etc.

In current study, we compared wildtype and PD trans-genetic *Drosophila melanogaster* with E46K mutation of SNCA gene for spontaneous behavioral classification and mapping. By using the pre-analysis software DeepLabCut (DLC), which provides marker-less pose estimation of user-defined body parts^13,14^, we monitored the posture information of *Drosophila melanogaster* during walking in a 3D-printed trap. Finally, we constructed a binary diagnosis system method based on extracting and clustering the multi-dimensioned behavioral features of these two types of *Drosophila melanogaster*. We hope this project provide an objective and quantitative behavioral assessment tool for the research on neural mechanism of PD with *Drosophila melanogaster*.

## Materials and Methods

### Animals

Two strains of *Drosophila Melanogaster* are included in this study: Oregon R strain wild type drosophila ordered from Carolina as the control (WT) group and the genetically modified α-Synuclein PD strain (SNCA^E46K^ mutant) ordered from Bloomington Drosophila Stock Center as the PD group. Both strains of drosophila are housed in test tubes, and maintained under constant conditions (12-h/12-h light/dark cycle, 23□, and a standard yeast-sugar-based food medium). Male and female are studied separately within each group because the behavioral changes of PD are usually more obvious for male.

### Behavioral apparatus and data collection

*Drosophila Melanogaster’s* movements are restricted into a specific area under the microscope with the 3D-printed trap (7mm*9mm*3mm, size decided upon the field of microscope). Videos of the WT and PD groups are recorded with a microscope and Motic software. The recording resolution was 2048×1536 at 25 frame-per-second (fps). We collected 40 animal videos in total (N_PD female_ = 11, N_PD male_ = 7, N_PD female_ = 14, N_PD male_ = 8).

### Drosophila melanogaster pose estimation with DeepLabCut

Before generating the pose estimation model, we resize the resolution (resized resolution: 512 ×384) of these collected videos to improve the computational efficiency. Thirty *Drosophila Melanogaster* videos were selected to generate the training set. We used the K-means clustering method to extract 20 frames from each video, and finally obtained a 600-frame to be labeled images. Then, we created a DeepLabCut (DLC, version: 2.2.0.6) project and configured nine pupil feature points to be detected: head, body center, tail, left front leg, left middle leg, left hind leg, right front leg, right middle leg, right hind leg (see Fig1). To train the DLC model, we used graphical user interfaces (GUIs) to manually label 600 extracted images. After completing the manual labeling step, the DLC model was trained in Python3 environment. These training steps were executed on the NVIDIA GeForce RTX 2080ti graphic processing unit. We set the maximum number of training iteration as 1030000, and it took more than 10 hours to complete these processes.

We used the trained model to track the 40 *Drosophila Melanogaster* videos and obtain 9 key body parts. Since each body part has 2 dimensions, each animal contains 18-dimension behavioral trajectory data.

### Behavioral data pre-processing

When collecting *Drosophila Melanogaster* videos, the distance between behavioral assay and the microscope lens was not always consistent. Therefore, for each video, we manually determined the trap boundary as ROI to ensure the consistency of the trap in the cropped image. Then we normalized the trajectory data, resizing values in the horizontal and vertical directions to a range of 0-100.

### Kinematic features extraction

In order to better observe the locomotion throughout the recording process, we plotted a velocity trajectory heatmap for each animal. We used the normalized trajectory of the body center point to represent the motion of whole body. First, we calculated the frame-by-frame velocity of each animal. Then to reduce noise and improve the visualization, we also smoothed the velocity with half a second time window. Finally, we used the blue-to-red colors in the heatmap to represent the low-to-high speed (Figure. 2A).

**Figure 1.**
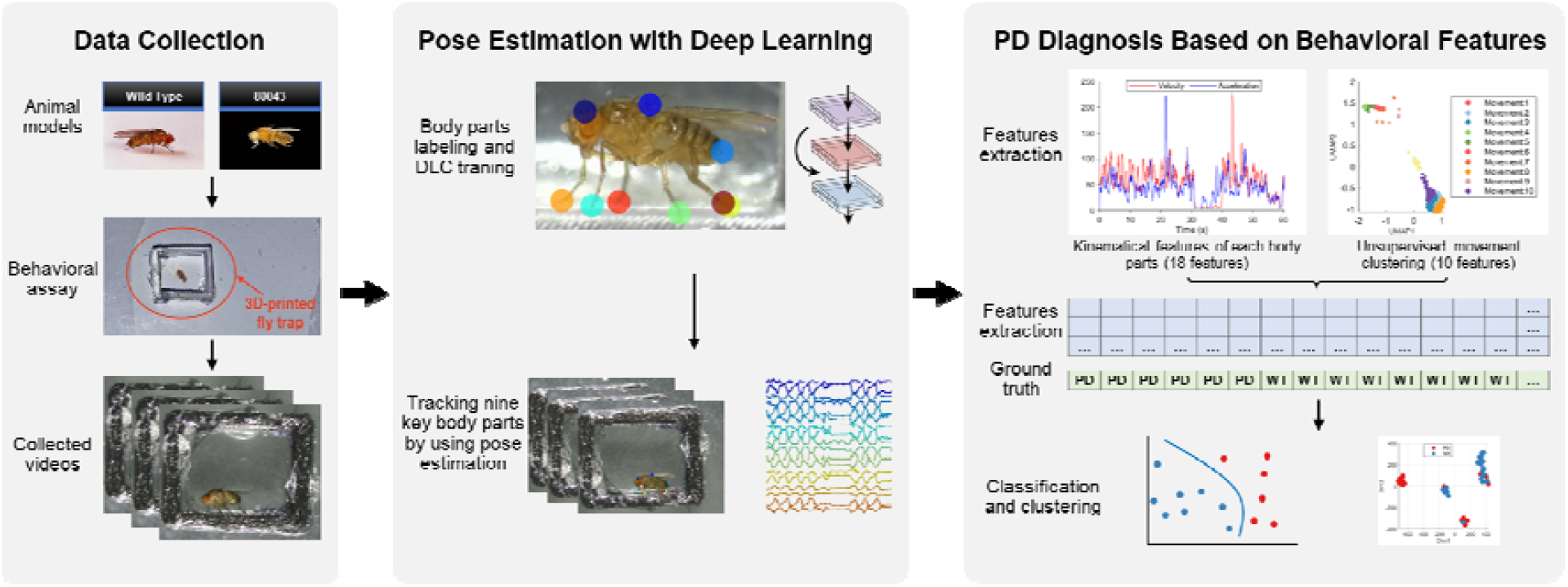
Diagram of deep-learning-based system for Parkinson’s Disease Drosophila melanogaster diagnosis.

**Figure 2.**
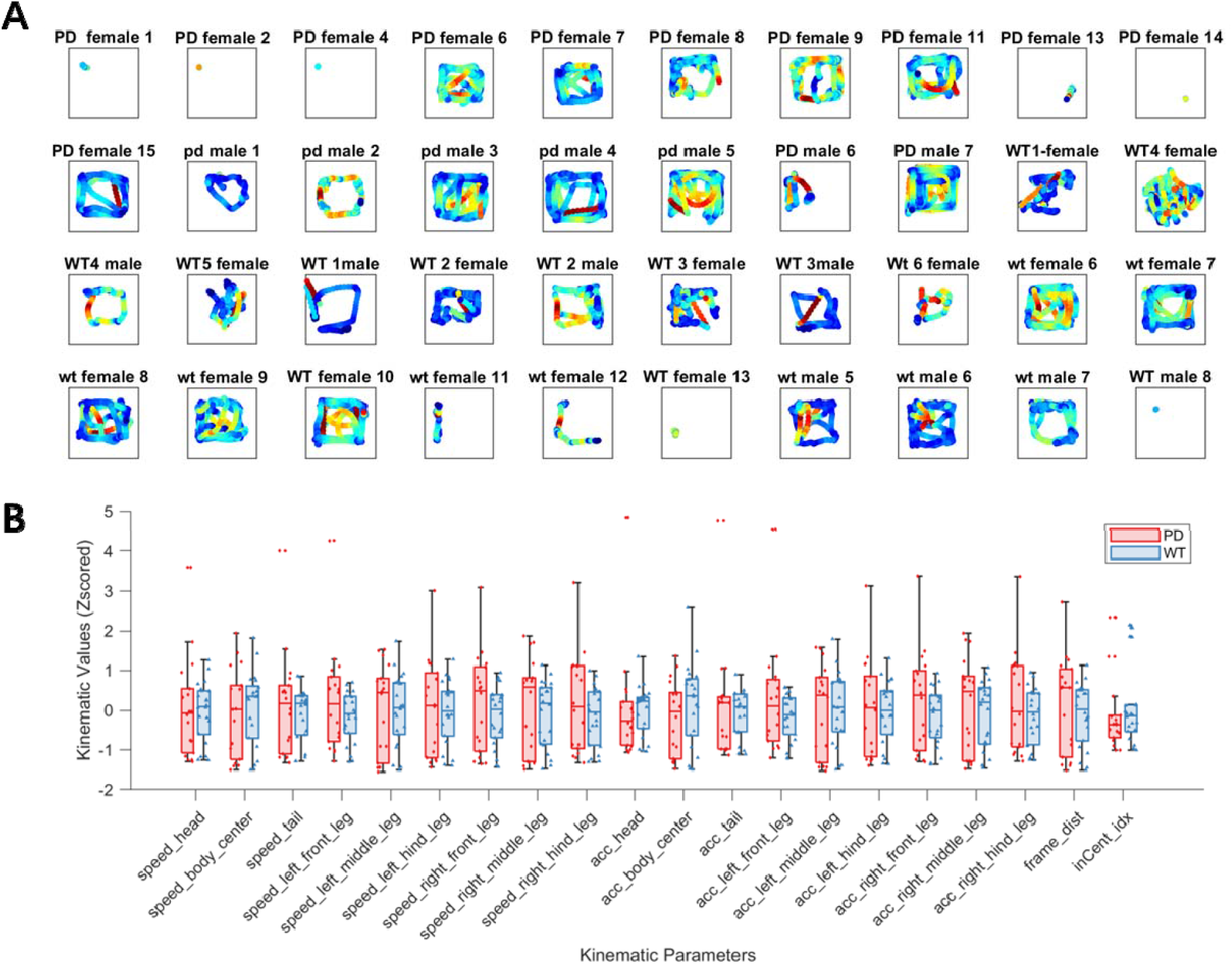
Comparison of kinematic parameters between WT and PD groups. A) Heatmap of velocity track of 40 *Drosophila melanogaster*. The activity track of the back point represents the overall movement. From blue to red in the heatmap, the speed is from low to high. B) Display the distribution of 20 kinematic parameters of PD and WT groups at the same scale, the values are normalized with z-score. The scatter points in the graph represents each sample. Boxplot is expressed in mean ± sem.

In addition, we calculated 20 kinematic features based on the normalized 18-dimensional body trajectories, including the velocity of 9 body parts, the acceleration of 9 parts, the per frame movement distance, and the ration of the animals entering the central region. We calculated these features as the following rules:

*1) Velocity*

We set T_*body*_ = {x_t_, y_t_},t = 1: *n* to be the coordinates of one of the body points, where *n* is the number of frames corresponding collected video. Therefore, the velocity of this body part can be calculated as follows:

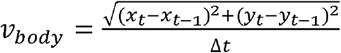

where Δ*t* is the frame rate of the recorded video.

*2) Acceleration*

Similarly, the acceleration of a body part can be calculated as follows:

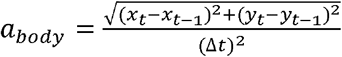

3) *Per frame distance*

We used the body center point T*_body center_* to calculate the per frame distance, the per frame distance of a body part can be calculated as follows:

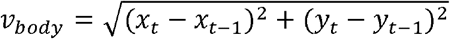

4) *Ratio in centre region*

Since we have normalized the trajectory data to the range of 0-100, for each frame, both the x and y coordinates of the body center in the range of 25-75 were considered to be in the centre. Then, the ratio in centre can be calculated as the ratio between the total number of frames in the centre and the total number of frames recorded.

### Unsupervised movement clustering

Since the symptoms of some mental diseases usually manifest themselves in specific stereotyped movements, we need to convert the 18-dimension trajectory data into movement sequences. We, therefore, used the Behavior Atlas (https://behavioratlas.tech/) to perform the trajectory decomposition and unsupervised movement clustering. Because the tracked 2D *Drosophila Melanogaster* trajectories were different from the 3D skeleton used in Behavior Atlas, it was difficult to perform the body alignment. Thus, we adopted part of the original method of Behavior Atlas. We calculated the dynamic time alignment kernel (DTAK) matrix of the 18 kinematic features (9 body parts velocity and 9 body parts acceleration) for each animal. After decomposing the trajectories of all animals, we calculated the 2D movement features space. Then we used hierarchical clustering to identify 10 clusters of these movement sequences.

### Dimensionality reduction and classification of Drosophila Melanogaster based on behavioral features

Twenty kinematic features and 10 movement subtypes were involved in the diagnosis of PD *Drosophila Melanogaster*. To intuitively evaluate the distribution between different types of animals according to behavioral features, we performed dimensionality reduction analysis with the t-SNE (t-distributed stochastic neighbor embedding) algorithm on the average value of these features. We used dimensionality reduction two-dimensional scatters and different colors to represent two types of animals (Figure 2A, B).

In the *Drosophila Melanogaster* classification section, we combined the kinematic features and the movement sequence fraction features into a 30×40 feature matrix. Correspondingly, the 40 PD and WT animals formed 1×40 labels (22 WT and 18 PD). We used the MATLAB (Version: 2020b, MathWorks, Massachusetts, USA) classification app and selected 6 models for training: Fine Tree, Weighted KNN (k-Nearest Neighbor), Fine KNN, Logistic Regression, Linear SVM (Support Vector Machine), Cubic SVM. Set validation to 10-fold cross-validation.

### Statistics

To compare kinematics and movement sequence fractions between the two groups, we imported these values for each animal using MATLAB and used Prism 8.0 (GraphPad Software, Inc.) for statistical analysis. Before hypothesis testing, the data were first tested for normality using the Shapiro-Wilk normality test and for homoscedasticity using the F test. One-way analysis of variance with the Kruskal–Wallis’s test was performed to determine which feature has a significant difference between the two groups.

## Results

### A Deep Learning-based System for Parkinson’s Disease Drosophila Melanogaster Diagnosis

*Drosophila melanogaster* PD diagnostic system based on deep learning includes three parts: 1) data collection (see in Figure 1). For WT and PD *Drosophila melanogaster*, we used a microscope to record the track spontaneous locomotion in a 3D-printed trap, and captured video of each animal. 2) *Drosophila melanogaster* pose estimation based on deep learning: based on the DeepLabCut pose estimation toolkit, 600 photos are manually marked, each of which is marked with 9 body points as a training set. Then we used the trained model to track the body trajectory of *Drosophila melanogaster*. 3) PD diagnosis of *Drosophila melanogaster* based on behavioral features: kinematics parameters of body motion and sequential activity characteristics are extracted respectively to form a matrix. Then, we reduce the dimension of visualization and use different machine learning models to achieve PD diagnosis with *Drosophila Melanogaster*.

### Kinematic Features are Insufficient for the Diagnosis of PD Drosophila Melanogaster

The variation of locomotor activity is a typical characteristic of PD in *Drosophila melanogaster*. We measure the velocity trajectory heatmap for each animal, and analyzed normalized trajectory of body center frame-by-frame to portrayal the action speed and spatial-temporal distribution of different genotype *Drosophila melanogaster* (see in Figure 2A). The color from blue to red represents speed from slow to fast. We found no difference in the velocity trajectory between WT and PD group, despite the female PD group presented higher probability of immobility. After that, we compared the kinematic features of *Drosophila melanogaster* with 20 parameters. We observed no difference in the velocity, acceleration, per frame distance, ratio in the centre individually (see in Figure 2B). These findings suggested that only kinematic features are insufficient for recognizing and distinguishing WT and PD *Drosophila Melanogaster*.

### Significant Difference between WT and PD Groups in the Movement Sequence Level

According to previous study of PD in *Drosophila melanogaster*, stereotyped behaviors is the most obvious alteration in PD *Drosophila melanogaster* compared to WT. Therefore, we establish an 18-dimension trajectory for clustering the spontaneous behaviors into 10 movement subtypes. Since the 10 movement subtypes occur in different proportions in each animal, we calculated the movement fraction of each animal. The movement fractions are defined as the bouts number of each movement phenotype divide by the total number of movement bouts the animal occurred during the experiment. Finally, we exacted two distinctive movement subtypes from 10 clusters (see in Figure 3A, B), No.1 and No7, which represent forelimb rubbing without body center movement in a corner, and forelimb dominated fast unilateral movement respectively (see in Figure 3C). These results revealed a significant difference between WT and PD *Drosophila melanogaster* in the movement sequence.

**Figure 3.**
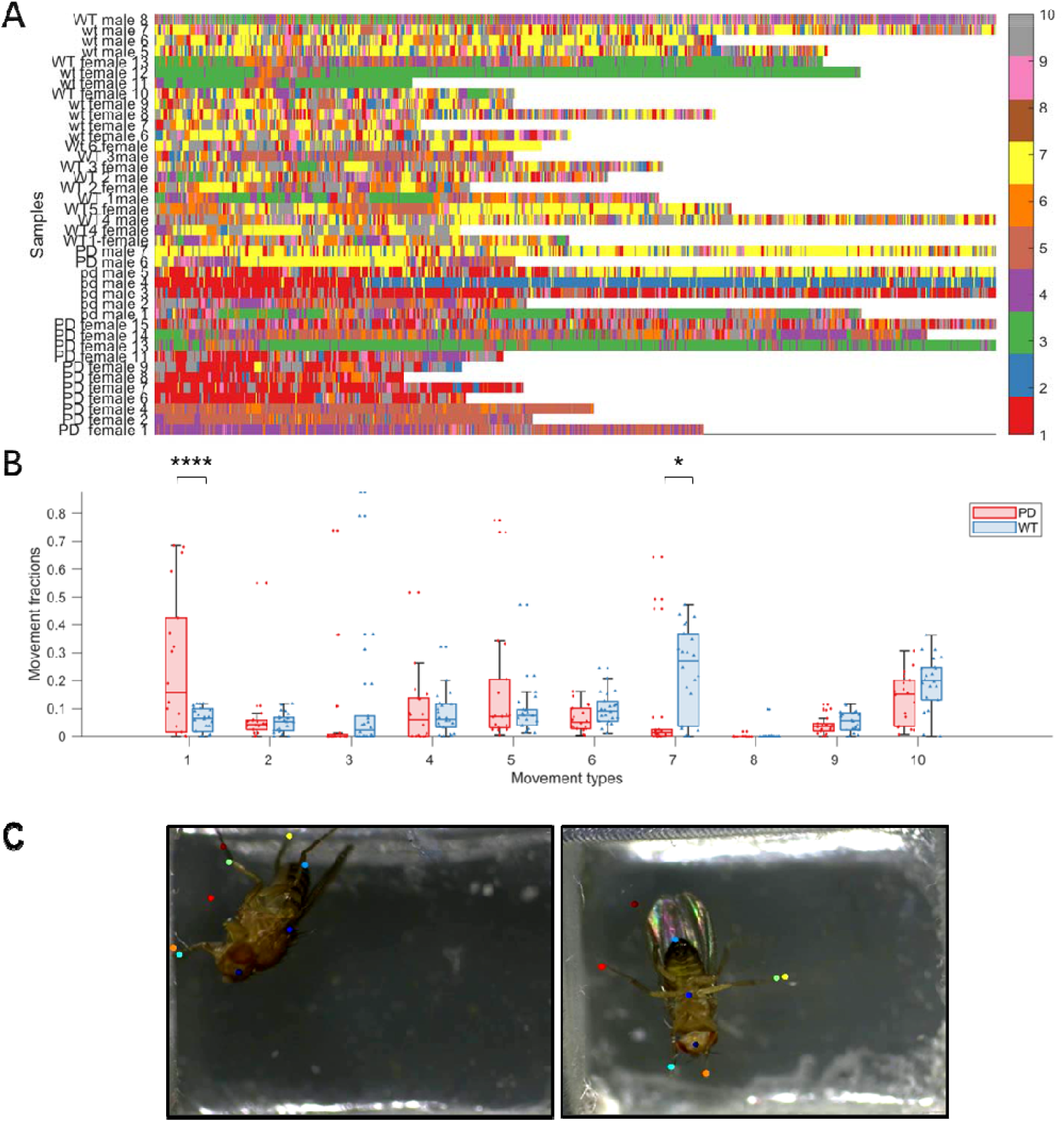
Comparison of movement fractions between WT and PD groups. A) Ethogram of 40 *Drosophila melanogaster*. Use 10 colors to show the types of spontaneous actions that *Drosophila melanogaster* perform during the corresponding period. Since the recording time of each animal is different, the end of the figure is not aligned. B) Comparison of 10 action types of PD and WT. Scatter points in the figure represent the fraction of each sample on this action, and boxplot is represented by mean ± sem. C) Typical behavioral performance of No.1(left) and No.7, which represent forelimb rubbing without body center movement in a corner, and forelimb dominated fast unilateral movement separately.

### Classification of Drosophila Melanogaster using combined features

To develop a deep learning binary PD diagnosis system based on extracting and clustering the multi-dimensioned behavioral features of these two types of *Drosophila melanogaster*, we further performed dimensionality reduction analysis with the t-SNE algorithm and testify the performance of different model for *Drosophila Melanogaster* classification. The results showed the behavior characteristics of WT and PD groups could be clearly separated into two clusters after dimensionality reduction (see in Figure 4A, B). In addition, compared to other methods for classification, the Fine Tree model has the highest accuracy (85%) and true positive rate (TPR, 88.9% in PD and 81.8% in WT), and the lowest false negative rate (FNR, 11.1% in PD and 18.2% in WT) (see in Figure 4C). The other models for classification, such as Weighted KNN, Fine KNN, Logistic Regression, Linear SVM, Cubic SVM, cannot achieve the same accuracy of PD diagnosis (see in Table 1). This result verified that our deep learning binary PD diagnosis system possessed the ability to classify PD *Drosophila Melanogaster* via behavioral mapping.

**Table 1.** Performance of different model for Drosophila Melanogaster classification.

| *Performance Methods* | *Accuracy* | *TPR (PD, WT)* | *FNR (PD, WT)* | *AUC* |
| --- | --- | --- | --- | --- |
| *Fine Tree* | 85.0% | 88.9%, 81.8% | 11.1%, 18.2% | 83.0% |
| *Weighted KNN* | 62.5% | 50.0%, 72.7% | 50.0%, 27.3% | 70.0% |
| *Fine KNN* | 62.5% | 50.0%, 72.7% | 50.0%, 27.3% | 70.0% |
| *Logistic Regression* | 57.5% | 66.7%, 50.0% | 33.3%, 50.0% | 62.0% |
| *Linear SVM* | 72.5% | 61.1%, 81.8% | 38.9%, 18.2% | 75.0% |
| *Cubic SVM* | 65.0% | 61.1%, 68.2% | 38.9%, 31.8% | 68.0% |

\* TPR: True Positive Rates; FNR: False Negative Rates; AUC: Area Under Curve

**Figure 4.**
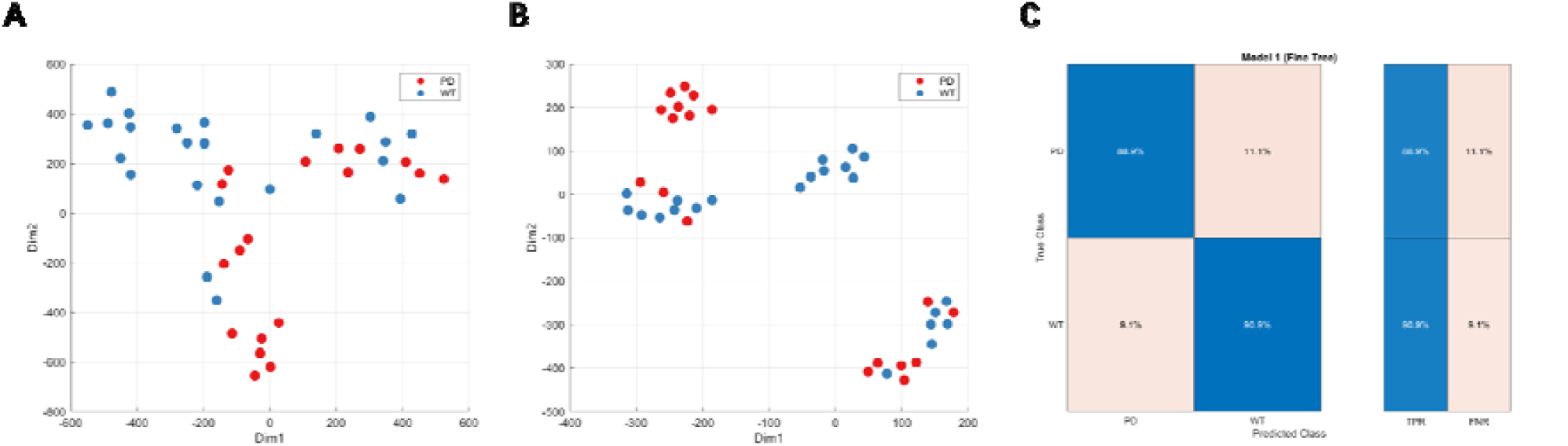
Clustering and classification of *Drosophila melanogaster* based on behavioral features. A & B) t-SNE algorithm is used to reduce the dimensions of kinematics and action sequences. The two colors represent clusters from different genotypes. C) The confusion matrix of Fine tree model with the best performance among the six machine learning models.

## Discussion

Previous studies were used to observe the survival and locomotion of *Drosophila melanogaster* larvae, and climbing ability, courtship, olfactory memory of adult to explore the effects of genetic, environmental and drug on PD phenotypes. However, the traditional climbing test (negative geotaxis) is tedious, labor-intensive, time-consuming, and deliver significant differences between different experiments^15^. *Alexander Mathis et al*. developed a DeepLabCut toolbox for markerless tracking of *Drosophila melanogaster*. DeepLabCut allows accurate extraction of low-dimensional pose information from videos of freely behaving Drosophila melanogaster^14^. Meanwhile, *Anqi Wu et al*. proposed a probabilistic graphical model built on top of deep neural networks, Deep Graph Pose (DGP), to leverage these useful spatial and temporal constraints and develop an efficient structured variational approach to perform inference^12^. For accelerating processing speed and improving robustness, *Jacob M Graving et al*. used an efficient multi-scale deep learning model, StackedDenseNet, for developing a novel friendly used toolkit DeepPoseKit, which increase processing speed by 2 times without loss of accuracy^11^.

In current study, we used the pre-analysis animal pose estimation software DeepLabCut (DLC) to transform the video data into numerical data representing the motion speed, tremor frequency, and limb motion of *Drosophila melanogaster*. We initially observed the locomotion dynamics of *Drosophila melanogaster* in 3D-printed trap, and compared the motion trajectory with 20 kinematic parameters of limbs speed and location. According to an unsupervised technology to discover and track the stereotypical behavior of drosophila recently developed by *Gordon J. Berman et al*., we compared the behavioral sequence of WT and PD *Drosophila melanogaster* with their alternative internal states^16^. We identified and clustered 10 typical behavioral features and found that PD *Drosophila Melanogaster’s* often perform forelimb rubbing and rapid shrinking abnormally, while forelimb-dominated rapid lateral movement significantly decreased. These phenotypes are similar to the bradykinesia and tremor in PD patients. The dichotomous PD diagnosis system developed in this project takes the massive data collected by DLC as input and outputs the diagnosis results. Moreover, we identified the best algorithm of clustering and classification is the Fine tree, of which the accuracy of prediction is the highest, up to 90%.

Parkinson’s disease is the second most prevalent neurodegeneration in the world. Not only PD cause a great burden of patients, but also lead to a significant increase in the risk of suicide^17^. Actually, the pandemic of Covid-19 also has a severe impact on the central nervous system, such as disrupting the blood brain barrier^18^. The spread of Covid-19 also challenges the prevention and the hospitalized management of PD patients^19,20^. The pathological progress of PD triggered by poisoning, infection and social stress is different from that with gene mutation. PD induced by environmental factors progresses slowly, with hidden or inaccurate symptoms. It will be hard to study the dose-effect of pharmacological therapy, the mechanism of chronic environmental stress in *Drosophila melanogaster* without objective and continuous dynamic observation. Our project has successfully constructed a data-driven diagnostic toolbox for PD research with *Drosophila melanogaster*.

